# Combinations of genomic alterations and immune microenvironmental features associate with patient survival in multiple cancer types

**DOI:** 10.1101/2024.12.27.630504

**Authors:** Masroor Bayati, Zoe P. Klein, Alexander T. Bahcheli, Mykhaylo Slobodyanyuk, Jeffrey To, Kevin C. L. Cheng, Jigyansa Mishra, Diogo Pellegrina, Kissy Guevara-Hoyer, Chris McIntosh, Mamatha Bhat, Jüri Reimand

## Abstract

Oncogenesis and tumor progression are shaped by somatic alterations in the cancer genome and features of the tumor immune microenvironment (TME). How interactions of these two systems influence tumor development and clinical outcomes remains incompletely understood. To address this challenge, we developed the multi-omics analysis framework PACIFIC to systematically integrate genetic cancer drivers and infiltration profiles of immune cells with clinical information. In an analysis of 8500 cancer samples, we report 34 immunogenomic interactions (IGXs) in 13 cancer types in which context-specific combinations of genomic alterations and immune cell activities associate with disease outcomes. Risk associations of IGXs are potentially explained by tumor-intrinsic and microenvironmental metrics of immunogenicity and differential expression of therapeutic targets. In luminal-A breast cancer, *MEN1* deletion combined with reduced neutrophils is associated with poor prognosis and deregulation of immune signalling pathways. These findings help elucidate how cancer drivers interact with TME to contribute to tumorigenesis.

## Introduction

The tumor immune microenvironment (TME) is a complex ecosystem of tumor cells, adjacent normal tissue, and immune cells that jointly contribute to tumor evolution. The immune system can sense and destroy tumor cells early in tumor evolution while later disruptions in TME and interactions with malignant cells enable a tumor-supporting environment. Tumors overexpress inhibitory immune checkpoints to evade T cell elimination [1], while chronic inflammation predisposes to cancer development and promotes all stages of tumorigenesis [2]. The complexity of the TIME across organs and patients poses a major challenge to current cancer treatments [3]. New mechanistic insights and molecular biomarkers are required to further characterise the TIME and develop personalized treatments [4]. However, therapeutic development is challenged by the genomic and molecular heterogeneity of tumors as well as surrounding tissue context and activity of the immune system [5, 6]. Immunotherapy efficacy associates with genomic instability and neoantigen burden of tumors [7, 8], as well as specific genetic alterations such as *BRAF* mutations and *PTEN* loss in melanoma [9, 10], *STK11* mutations in *KRAS*-mutant lung adenocarcinoma [11], and *EGFR* mutations in non-small cell lung cancer [12]. Epigenetic reprogramming, signalling pathway activities, and altered metabolism also interact with TME [13]. Microenvironmental interactions have been explored systematically using functional genomics screens *in vitro* and *in vivo* [14, 15], and using multi-omics analyses of cancer patient cohorts to understand the interactions of genomic alterations and signatures of immune cells [16–20]. Recent advances in single-cell sequencing have revolutionized precision oncology by dissecting the transcriptional dynamics of cancer cells with stromal and immune cells, providing insights into tumorigenesis and therapeutic resistance [21, 22]. However, profiling large clinical cohorts of cancer patients to characterise TME using single-cell technologies is not yet feasible. To our knowledge, the pan-cancer landscape of synergistic interactions of genomic alterations and TME with respect to clinical outcomes in cancer have not been systematically explored to date.

We hypothesized that genomic alterations in cancer cells may have synergistic interactions with TME and immune features that are reflected in clinical outcomes. Using our novel statistical framework PACIFIC, we integrated immune cell signatures from bulk transcriptomes with recurrent genomic alterations and patient clinical profiles from matching samples. Using the TCGA PanCanAtlas cohort of 8500 primary, untreated cancer samples of multiple cancer types, we identified a set of immunogenomic interactions (IGXs) in 13 cancer types that defined high-risk subsets of cancer samples that were not apparent when analysing either the genomic or immune features of the cancers alone. We focused on one high-confidence IGX found in luminal-A breast cancer in which deletions of the *MEN1* tumor suppressor gene together with reduced neutrophil levels associates with worse prognosis and transcriptional deregulation of immune signalling pathways. This study illuminates an underexplored intersection of genetic cancer drivers and TME with potential insights into cancer biology and translational applications.

## RESULTS

### PACIFIC: a statistical framework for finding survival-associated interactions in multi-omics data

We developed the computational method PACIFIC to systematically discover multi-omics interactions that associate with clinical variables such as patient survival time or tumor subtypes. The algorithm highlights multi-omics interactions such that the co-occurrence of two omics features associates more strongly with patient survival or another clinical variable than either of the omics features alone (**Figure 1A**). Genomic alterations such as mutations or copy number alterations (CNAs) as well as transcriptomic, epigenomic or proteomic profiles can be used as the predictive multi-omics features (**Figure 1B**). The method uses patient survival information to find associations with omics profiles while continuous and binary clinical variables are also permitted. To prioritise complementary omics signals, baseline information such as patient age and sex and tumor stage or grade are also included in the models as covariates.

**Figure 1.**
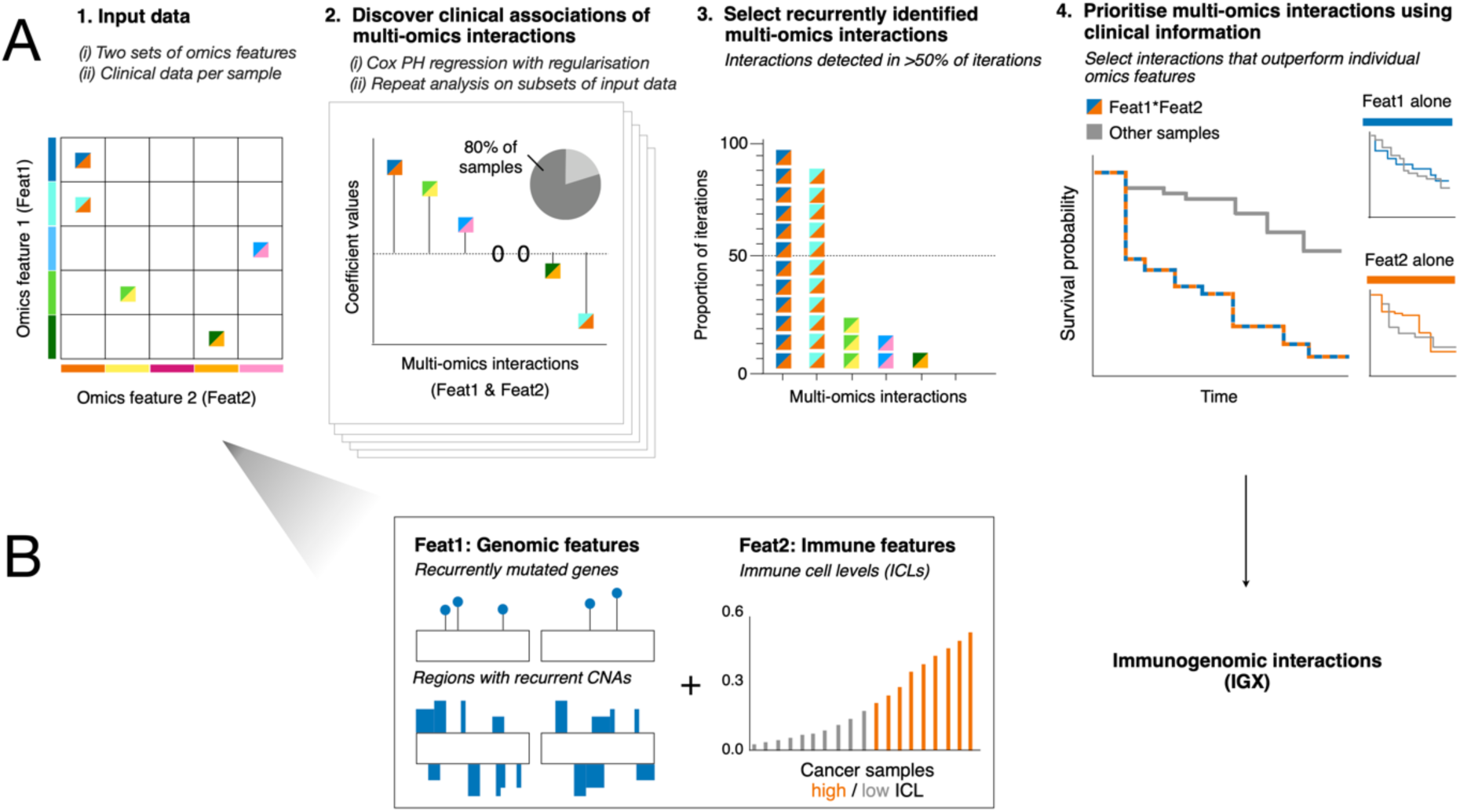
Discovery of multi-omics interactions with clinical significance using PACIFIC. **(A)** PACIFIC integrates pairs of multi-omics features (Feat1, Feat2) with patient clinical information to find multi-omics interactions (i.e., pairs of omics features) that associate with patient survival or other clinical variables. Two major types of input data are required (i) two sets of molecular features from omics experiments annotated per sample, and (ii) patient survival information and other clinical information. (2) To select multi-omics interactions, we fit regularised survival regression models using subsets of samples as input over a series of iterations and select interactions with non-zero CoxPH coefficients. (3) We prioritise multi-omics interactions identified in at least half (50%) of the iterations and (4) perform further filtering to select those exceeding the survival associations of Feat1 and Feat2 alone, and clinical variables alone. **(B)** To find immunogenomic interactions (IGX), we investigate multi-omics interactions of genomic features (i.e., significantly mutated genes and recurrent CNAs) and immune features (i.e., immune cell infiltration profiles from matching bulk cancer transcriptomes). We applied PACIFIC to 8,450 primary tumors across 26 cancer types in the TCGA PanCanAtlas using these multi-omics features.

To identify clinically associated omics interactions systematically, we adapted generalised linear regression analysis with elastic net regularisation using omics profiles as model predictors and patient survival profiles as responses based on our prior work on cancer biomarker and target discovery [23, 24]. The algorithm first transforms individual omics profiles, such as genomic alterations and transcriptome-derived immune signatures, to pairwise combinations of features. Next, to derive candidate omics interactions, we select the most informative omics interactions for a randomly sampled subset of the patient cohort using elastic net regularisation and repeat this process over a series of iterations (**Methods**). The final omics interactions are selected based on two criteria: the omics interaction has been identified in at least half of the iterations, and its survival association exceeds the significance of the baseline clinical variables and the individual omics features comprising the omics interaction. The method assumes that input features are preprocessed appropriately. Here, we combined driver gene mutations (present or absent), chromosomal regions with CNAs (present or absent), and immune cell infiltration levels in tumors (high or low). We removed sparse omics interactions and those lacking univariate associations with patient survival. Identifying clinically informative omics interactions using PACIFIC allows the discovery of biomarker and mechanistic hypotheses through multiple layers of cellular logic. PACIFIC (Predict and Analyse Combinations of Interacting Features in Cancer) is available as an open-source R package at https://github.com/reimandlab/PACIFIC.

### Identification of survival-associated immunogenomic interactions

We asked if recurrent genomic alterations and immune cell infiltration profiles jointly associated with disease outcomes. We defined immunogenomic interactions (IGXs) as pairs of genomic features and immune features that are significantly associated with patient survival given clinical covariates (patient age, sex, tumor stage and/or grade) and exceed the survival association of the genomic or immune features alone (**Figure 1B**). To this end, we leveraged matched genomic, transcriptomic, and clinical profiles of 8450 primary tumors of 26 cancer types in TCGA PanCanAtlas [25] (**Table S1**). Two types of genomic features included 299 significantly mutated genes (SMGs) with single nucleotide variants (SNVs) or indels and recurrent CNAs retrieved from previous studies [26, 27]. Immune features were defined as the immune cell levels (ICLs) of 22 cell types derived from matching bulk cancer transcriptomes using the CIBERSORT method [18, 28] and were subsequently grouped as High or Low using median dichotomisation. We used overall survival or progression-free survival information to map clinical associations of IGXs, as recommended for different cancer types [29].

We found 34 IGXs in 13 cancer types (**Figure 2A; Figure S1; Table S2**). All IGXs were cancer type-specific and the genomic features contributing to IGXs were also distinct. IGXs involved 11 frequently mutated driver genes and 18 recurrent CNAs, each of which was found in a single cancer type. Exceptionally, driver mutations in *PIK3CA* were found in IGXs in squamous cell lung cancers (LUSC) and head & neck cancers (HNSC), while *KMT2D* mutations were found in IGXs in colon and uterine cancers (COAD, UCEC). While genomic features were predominantly cancer type-specific, immune cell activities in IGXs were often representative of multiple cancer types. Infiltration levels of naïve B cells were most frequently associated with patient survival and spanned five IGXs and four cancer types through interactions with different genomic features. We validated IGXs computationally and confirmed that their survival associations complemented common clinical variables and respective immune and genomic features alone, as required by PACIFIC (**Figure 2B-C**). Most IGXs defined small subsets of aggressive samples from each cancer type, representing 35 samples on average (**Figure 2D**). Sample subsets defined by IGXs are relatively small as the recurrence rate of most genomic alterations in IGXs is low and cancers are further grouped by immune cell levels.

**Figure 2.**
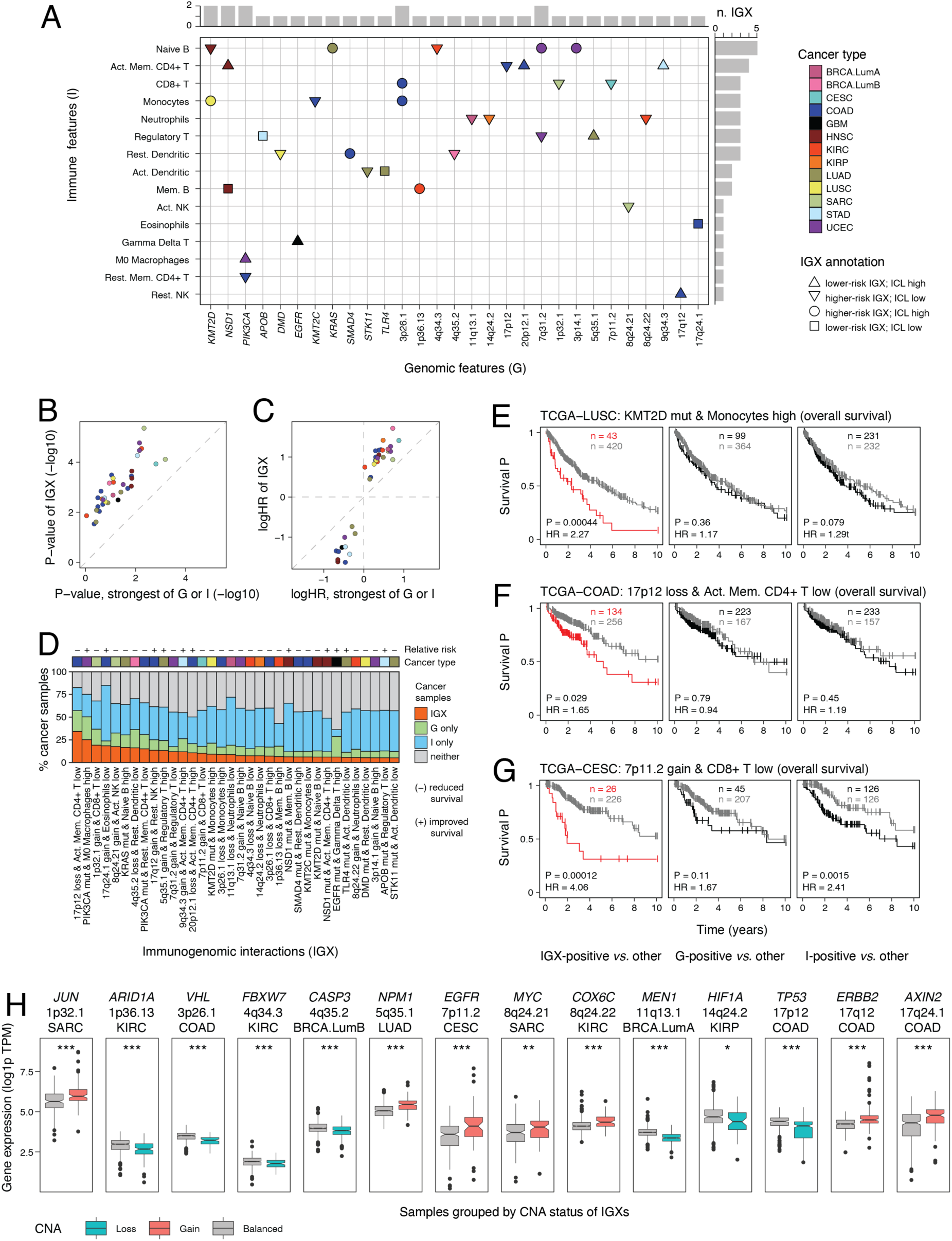
Immunogenomic interactions (IGX) associate recurrent genomic alterations and immune cell activities with clinical outcomes. **(A)** Summary of 34 IGXs identified in 13 cancer types. Each IGX comprises a pair of a genomic feature (X-axis) and immune feature (Y-axis) whose co-occurrence associates with patient survival in one cancer type (colors). Node shape indicates risk profile of the IGX and annotation of the immune feature (immune cell level as Low or High). **(B, C)** IGXs provide additional information about patient survival compared to the corresponding genomic or immune features alone. Scatter plots compare the survival analysis statistics of each IGX (Y-axis) and as controls the best associations from the genomic or immune feature of the IGX. Colors indicate cancer types. **(D)** Fraction of cancer samples in each IGX. IGX-positive samples (orange) are compared with samples with only genomic features (green), samples with only immune features (blue), and samples with neither features (grey). **(E-G)** Examples of IGXs. Kaplan-Meier (KM) plots compare overall survival (OS) of IGX-positive and IGX-negative subsets of cancer patients (left). As controls, OS analyses of the corresponding genomic and immune features are shown (middle, right). (E) *KMT2D* mutations combined with higher monocyte levels associate with worse prognosis in lung squamous cell carcinoma (LUSC). **(F)** Chr 17p12 (*TP53*) losses combined with lower levels of activated memory CD4+ T cells associate with worse prognosis in colon adenocarcinoma (COAD). **(G)** Chr 7p11.2 (*EGFR*) gains combined with lower CD8+ T cell levels associate with worse prognosis in cervical squamous cell carcinoma (CESC). P-values and HR values in **C and E-G** were derived from Cox PH models and ANOVA analyses and account for baseline clinical covariates. (**H**) Recurrent CNAs in the IGXs associate with oncogene activation and inhibition of tumor suppressor genes. Gene expression levels were compared between samples with and without the CNAs in the corresponding cancer types. We selected known cancer genes from the Cancer Gene Census from the altered regions (Wilcoxon test, FDR < 0.05; * < 0.05; ** < 0.01; *** < 0.001).

We studied the most prominent examples of IGXs (**Figure 2E, F, G**). Mutations in the tumor suppressor gene *KMT2D* combined with higher monocyte levels marked a high-risk subset of 43 lung squamous cell carcinomas (LUSC) (P = 4.4×10^-4^; hazard ratio (HR) = 2.27; **Figure 2E**). *KMT2D* encodes an epigenetic modifier whose inactivating mutations induce a metabolic shift towards glycolysis in lung cancer [30] and increase receptor tyrosine kinase signalling [31]. Elevated levels of monocytes associate with chronic unresolved inflammation that facilitates cell proliferation and contributes to cancer initiation and progression [32].

Genomic losses of 17p12 combined with fewer activated memory CD4+ T cells associated with worse prognosis in colon adenocarcinoma (COAD) in an IGX representing 134 samples (P = 0.029, HR = 1.65; **Figure 2F**). Recurrent genomic losses at 17p12 caused transcriptional downregulation of the adjacent tumor suppressor gene *TP53* (FDR = 4.1 x 10^-14^; **Figure 3C**), suggesting the functional role of 17p12 loss in this IGX. On the other hand, tumor-infiltrating CD4+ T cells contribute to adaptive immune response against malignant cells [33]. The association with worse prognosis suggests that p53 loss with reduced immune response may relate to worse disease outcomes.

**Figure 3.**
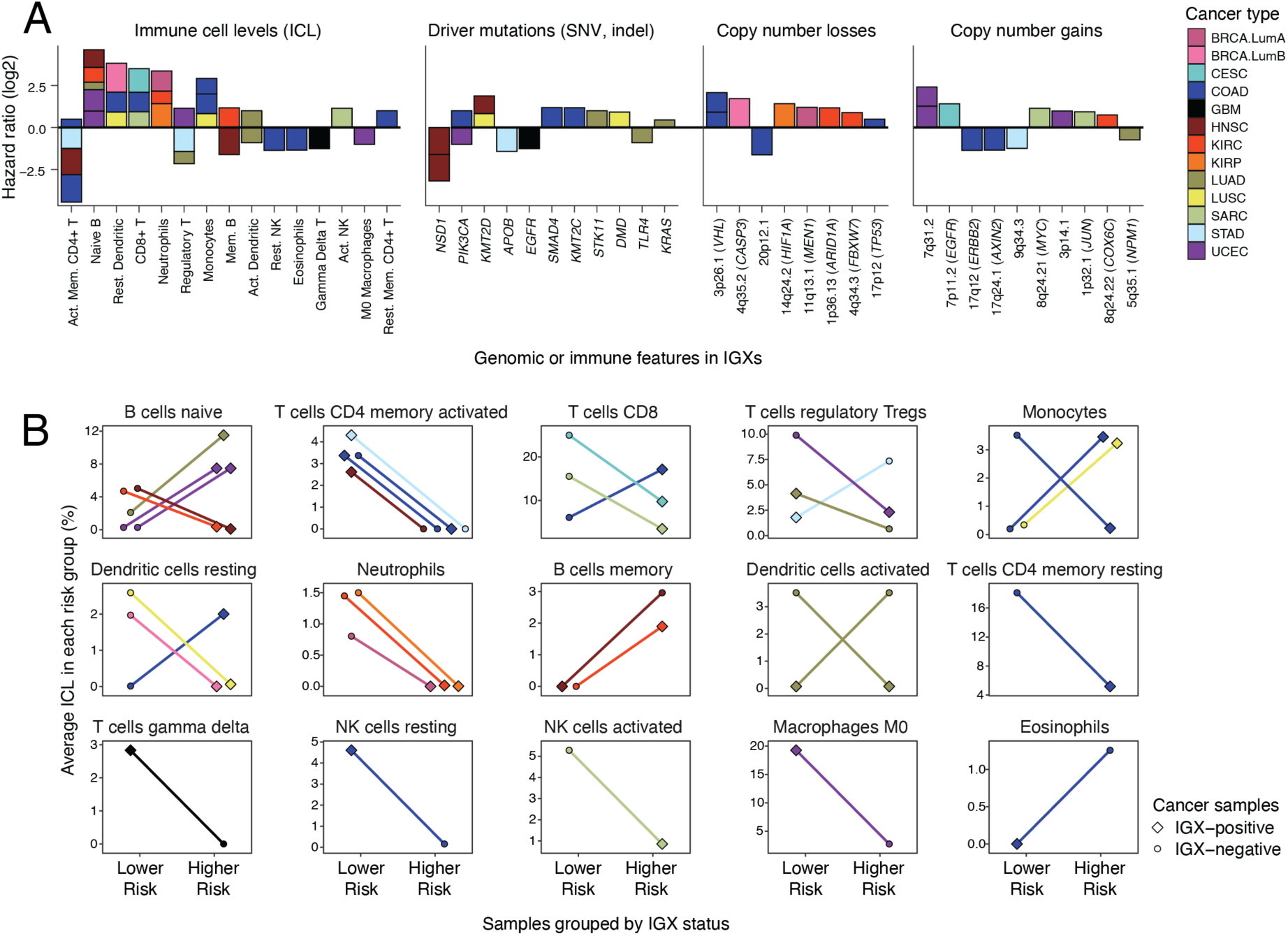
Molecular and clinical heterogeneity of IGXs. **(A)** Survival associations of the genomic and immune features involved in the IGXs. Bar plots show the aggregated risk scores of the features embedded in the IGXs. Risk scores are the sums of log-transformed HR values associated with each IGX that account for baseline clinical variables and genomic (G) and immune (I) features as covariates. **(B)** Immune cell levels (ICLs) included in IGXs show diverse associations with patient prognosis. Lines show immune cell levels (ICLs) of cell types contributing to the IGXs in high-risk and low-risk IGX groups (denoted by point shapes). Uniformly ascending or descending lines show immune features that have consistent associations with patient survival, while crossing lines indicate inconsistent, heterogenous associations with patient survival.

Genomic gains at 7p11.2 gains combined with fewer CD8+ T cells highlighted a subset of 26 cervical cancers (CESC) with worse prognosis (P = 1.2 x 10^-4^, HR = 4.06; **Figure 2G**). The 7p11.2 locus includes the oncogene *EGFR* (epidermal growth factor receptor) that controls cell proliferation, migration, and anti-apoptotic pathways. *EGFR* overexpression or mutation associates with poor prognosis in epithelial cancers and is an established therapeutic target in several cancer types [34]. In our data, IGX-positive CESC samples also show increased *EGFR* expression (P = 2.9 x 10^-6^, U-test; **Figure S2**), suggesting that 7p11.2 amplifications increase EGFR pathway activity that have functional interactions with TME, potentially in the context of this IGX. On the other hand, CD8+ T cells target tumor cells and drive adaptive anti-tumor immune responses [35] whereas lower levels of CD8+ T cells confer reduced anti-tumor immunity and worse prognosis [36]. Here, lower levels of CD8+ T cells also independently associated with poor prognosis in CESC; however, their combination with *EGFR* amplification in this IGX showed a pronounced effect on prognosis. These IGXs exemplify the findings in our catalogue and their further investigation may lead to functional and translational insights.

Focusing on 13 recurrent CNAs in the IGXs, we asked if oncogenes or tumor suppressors encoded by the genomic regions were deregulated transcriptionally by the CNAs. Indeed, genomic deletions led to significant downregulation of tumor suppressor genes such as *TP53*, *ARID1A*, *VHL*, *MEN1*, *CASP3*, *FBXW7,* and *HIF1A*, while genomic gains led increased expression of oncogenes such as *MYC*, *EGFR*, *ERBB2*, *COX6C*, *NPM1*, and *JUN* (**Figure 2H**). For instance, 1p36.13 loss was associated with *ARID1A* downregulation in kidney renal cell carcinoma (KIRC) while 8q24.21 gains led to *MYC* upregulation in sarcoma (SARC). In addition, numerous adjacent genes were also altered by CNAs and may contribute to tumor phenotypes and heterogeneity. Thus, genomic alterations in IGXs contribute to deregulation of hallmark cancer genes.

### Molecular and clinical heterogeneity of IGXs

Next, we studied the relative risk associations of IGXs. Most IGX-positive samples were associated with worse prognosis compared to other samples in the corresponding cancer type (23/34 or 67%). We quantified the genomic and immune features in terms of survival associations across all IGXs and confirmed that most features were found either in higher-risk IGXs or lower-risk IGXs. Activated memory CD4+ T cells, naive B-cells, and mutations in *NSD1* showed the strongest aggregated risk scores (**Figure 3A**).

Genomic alterations embedded in IGXs included known tumor suppressor genes and oncogenes whose whereas their genomic alterations were often consistent with their suppressive or promoting roles. Mutations in driver genes in high-risk IGXs occurred in *KMT2D*, *SMAD4*, *KMT2C*, *STK11*, *DMD*, and *KRAS*, while lower-risk IGXs involved mutations in *NSD1*, *APOB*, *EGFR*, and *TLR4* (**Figure 3A**). As expected, most genomic alterations involved in multiple IGXs associated with consistently increased or reduced risk. As an interesting exception, *PIK3CA* mutations showed opposite risk associations in two IGXs: *PIK3CA* mutations associated with worse prognosis in colorectal cancer when combined with lower levels of resting CD4+ T cells, while *PIK3CA* mutations in UCEC associated with improved prognosis when combined with higher M0 macrophage levels. Frequent mutations in the *PIK3CA* oncogene drive invasive phenotypes and have been found in several cancer types [37]. This suggests that genomic driver events may associate with different outcomes depending on the TME context.

Next, we studied the immune features involved in IGXs and found diverse clinical associations (**Figure 3B**). Several immune cell types associated with both better and worse prognosis in various IGXs depending on the context of the cancer types and the genomic features with which the interactions were identified. For example, naive B-cells, CD8+ T-cells, and regulatory T-cells were found in multiple IGXs representing both higher- and lower-risk cancers. Cytotoxic CD8+ T cells drive adaptive anticancer immunity and are a central focus in the immunotherapy development [38]. B cells are multi-faceted regulators and effectors of anti-tumor responses through antigen presentation and antibody production capabilities and their effector mechanisms in TIME is highly context dependent [39]. In comparison, activated CD4+ memory T cells were also found in four IGXs in different cancer types and associated with survival positively. CD4+ T cells coordinate CD8+ T cells and other immune cells to exert effective antitumor immunity and emerging evidence proves their direct anticancer effector roles [33]. Overall, the different risk associations of immune cell activities across IGXs reflect a heterogeneous nature of TME.

### IGXs associate with immunogenicity and therapeutic targets

To interpret the IGX functionally, we studied a panel of immunogenic and TME characteristics of matching cancer samples derived from previous TCGA studies [17, 18]. These included tumor-intrinsic genomic characteristics such as genomic instability, mutation load, and neoantigen burden, as well as immune characteristics such as cytolytic activity and interferon gamma pathway response [17, 40]. We also analysed gene expression levels of four major targets of immune checkpoint inhibitor (ICI) therapy: PD-1 (programmed cell death protein 1, *PDCD1*), PD-L1 (programmed cell death 1 ligand 1, *CD274*), CTLA-4 (cytotoxic T-lymphocyte associated protein 4, *CTLA4*), and LAG-3 (lymphocyte activation gene 3 protein, *LAG3*).

Comparing IGX-positive and IGX-negative samples revealed statistical associations with immunogenic and TME characteristics (FDR < 0.05; **Figure 4A**). Among the genomic characteristics, higher-risk IGXs were associated with increased genomic instability and intratumor heterogeneity in BRCA, COAD, KIRC, UCEC and SARC, suggesting that these IGXs characterise later-stage, genetically advanced and more aggressive tumors. Several IGXs were associated with reduced expression of ICI targets, indicating an immune-cold environment. However, cancers with homologous recombination deficiency (HRD) indicated by certain IGXs may be sensitive to PARP inhibitor therapies [41], providing a potentially actionable interpretation of IGXs. Among the immune characteristics, higher-risk IGXs were associated with lower levels of cytolytic activity that reflects suppression of anti-tumor immunity and offers a potential explanation to poor clinical outcomes.

**Figure 4.**
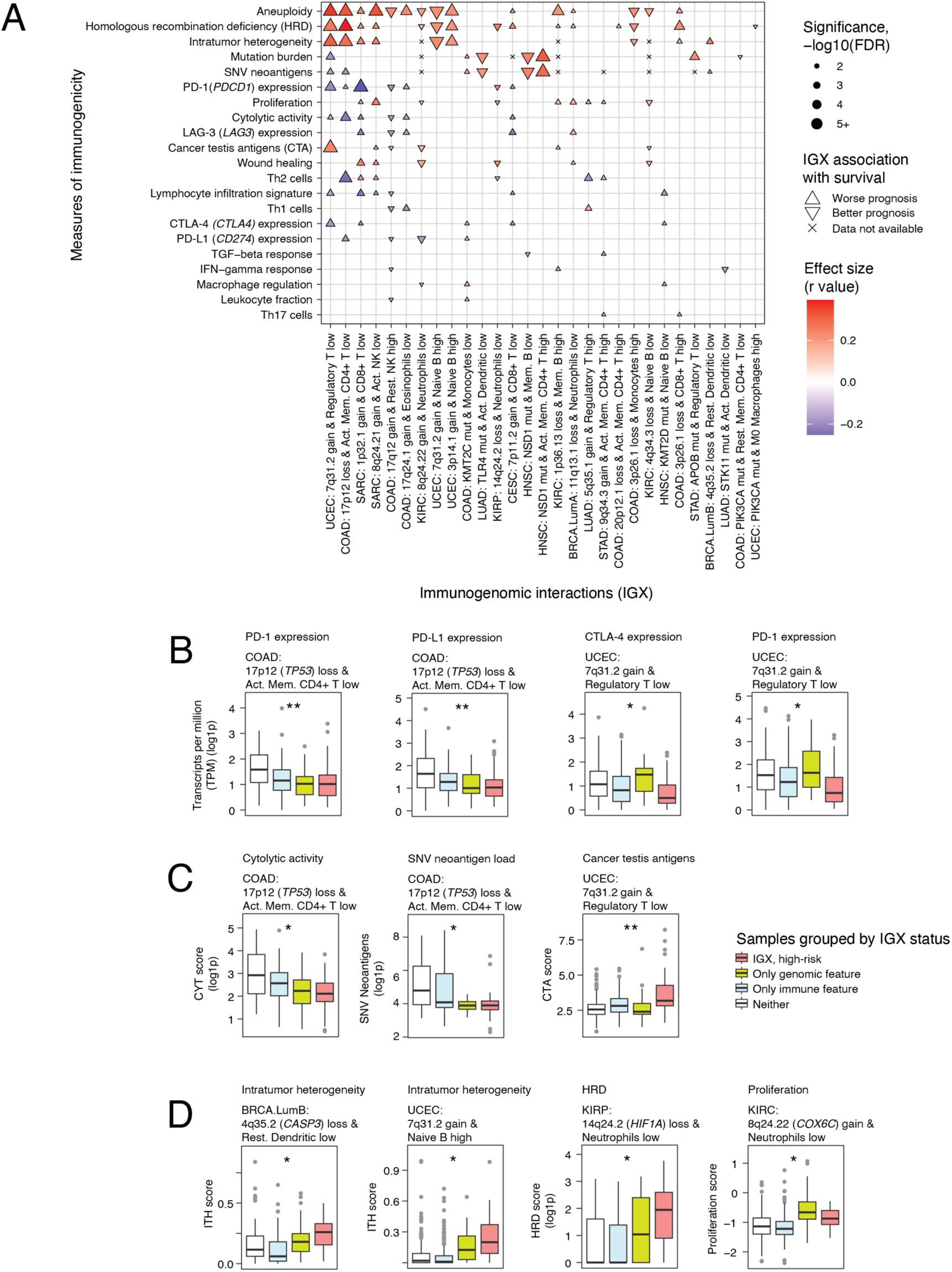
IGXs associate with measures of immunogenicity and therapeutic targets. **(A)** Associations of IGXs with genomic and immunogenic characteristics. IGX-positive samples were compared with IGX-negative samples across a panel of genomic and TME characteristics (FDR < 0.05, Mann–Whitney U test). IGXs are grouped as higher-risk (upward triangle) or lower risk (downward triangle). **(B-D)** High-confidence associations of IGXs that control for genomic and immune components of the IGX (ANOVA, P < 0.05 from interaction-informed regression analyses). Boxplots show the scores in four groups of cancer samples grouped by the presence of IGX (red) and the corresponding genomic (green) and immune (blue) components of the IGX, and samples with neither genomic or immune features (white). **(B)** High-confidence interactions of IGXs and gene expression of targets of immune checkpoint inhibition. **(C)** High-confidence interactions of IGXs and immunogenic characteristics of cancers. **(D)** High-confidence interactions of IGXs and genomic characteristics of cancers.

As a more stringent analysis, we asked which of the identified immunogenomic characteristics associated more strongly to the specific IGXs than expected from the genomic and immune features of the respective IGXs alone. We identified 11 strong associations involving six IGXs in an interaction-informed regression analysis to emphasize IGX effects (ANOVA *P* < 0.05, **Figure 4B-D**). For example, high-risk tumors in COAD were characterised by a co-occurrence of *TP53* loss and lower levels of activated memory CD4+ T cells. These IGX-positive colorectal cancers associated consistently with lower cytolytic activity, reduced expression of *PD1* and *PDL1*, and lower neoantigen burden. In UCEC, 7q32.1 (*EGFR*) gains and lower levels of regulatory T-cells occurred in higher-risk cancers with reduced levels of ICI targets and elevated intratumor heterogeneity. These IGXs may correspond to immunosuppressed, poorly treatable TME in a subset of cancers. This analysis provides complementary functional insights into IGXs and may lead to testable hypotheses for therapeutic development.

### *MEN1* loss and reduced neutrophils in luminal A breast cancer

Lastly, we focused on an IGX in luminal-A breast cancer involving genomic loss of 11q13.1 combined with reduced neutrophil levels that associated with lower progression-free survival in 47 of 538 cancers in TCGA (P = 0.0018, HR = 3.32; **Figure 5A**). To validate this interaction, we analysed an independent cohort of 414 luminal-A breast cancers from the METABRIC study [42], and confirmed that the IGX associated with reduced relapse-free survival in 25 samples (5.4 x 10^-4^, HR = 3.32; **Figure 5B**). 11q13.1 loss alone also associated with relapse-free survival in METABRIC and showed a confirming trend in TCGA, while neutrophil levels were not significantly associated with progression or relapse. We also evaluated overall survival and found confirming trend in one cohort (**Figure S3**). Validation of these associations in an independent cohort of cancer samples lends confidence to this IGX as a candidate biomarker and for future functional studies.

**Figure 5.**
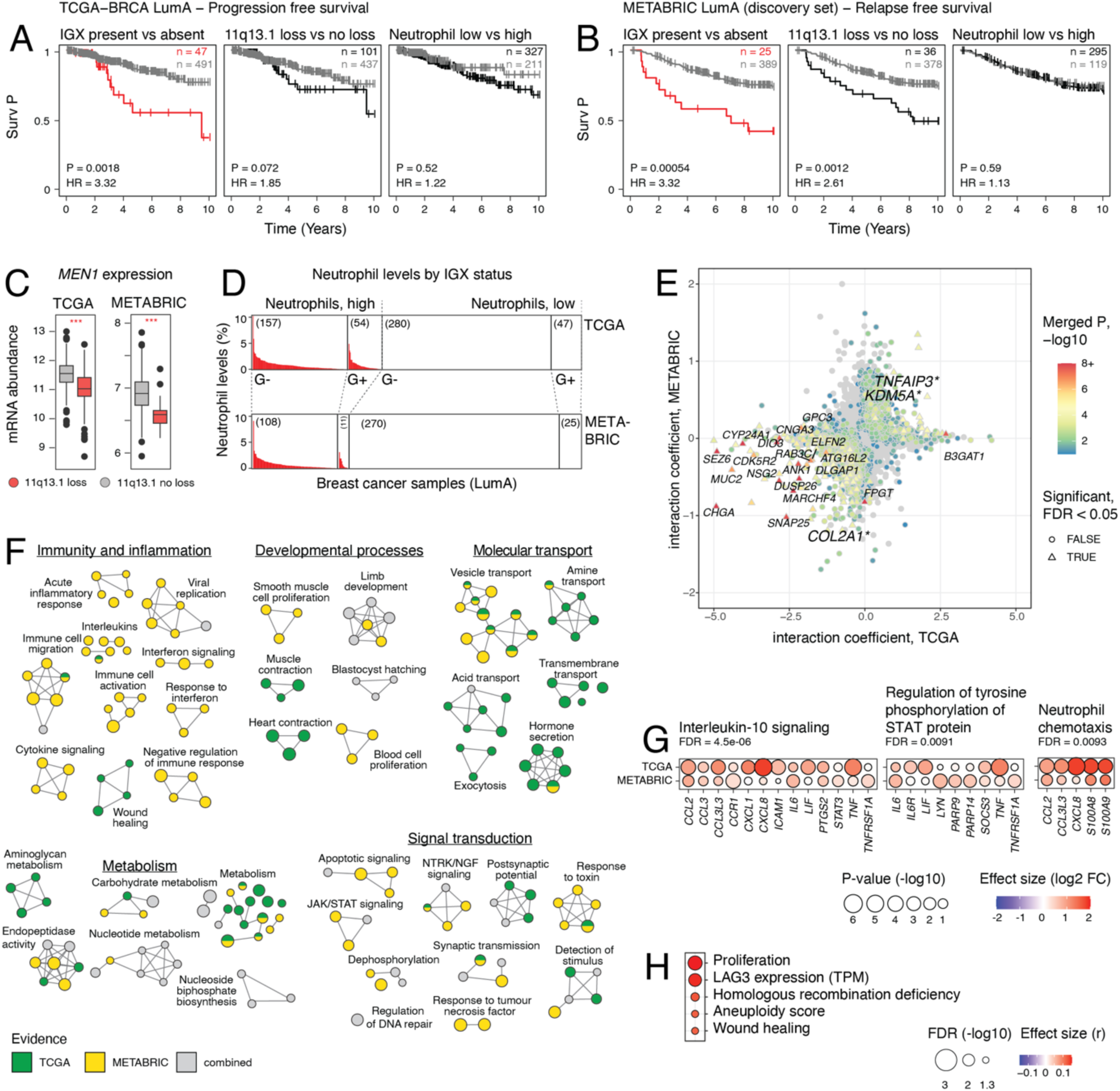
Genomic loss of *MEN1* at 11q13.1 and reduced neutrophils associates with tumor progression in luminal-A breast cancer. **(A-B)** Association of the 11q13.1-neutrophil IGX with progression-free survival (PFS) in luminal-A breast cancers in TCGA (panel A) and in an independent cohort from the METABRIC project (panel B). Kaplan-Meier (KM) plots on the left in (A-B) compare IGX-positive and IGX-negative samples when controlling for clinical covariates (age, stage, sex). Control analyses including only the genomic feature (11q13.1 loss) or the immune feature (reduced lelvels of neutrophils) are also shown (middle, right). Sample sizes (n) are shown. P-values and HR values were derived from clinically-adjusted CoxPH models. **(C)** *MEN1* expression levels in luminal-A breast cancers with and without 11q13.1 genomic deletions. **(D)** Relative abundance of neutrophils in samples grouped by IGX status, 11q13.1 (loss or no loss; indicated by G+ and G-) and neutrophil levels (low or high). **(E)** Differential gene expression analysis of IGX-positive luminal-A breast cancers that prioritises genes associated with IGX status in both the TCGA and METABRIC cohorts. The scatter plot shows coefficients of the IGX group controlled for G and I groups for TCGA (X-axis) and METABRIC (Y-axis). Genes are colored by the merged P-value from the two cohorts and triangles indicate significant genes (Brown merged FDR < 0.05). Known cancer genes from Cancer Gene Census (asterisks; FDR < 0.1) and highly significant genes (P < 1e-6) are labelled. **(F)** Pathways enriched in IGX-positive cancers visualised as an enrichment map. Nodes in the network represent significantly enriched pathways across the two cohorts of luminal-A breast cancers (ActivePathways, FDR < 0.05). Subnetworks group pathways with many shared genes, and node size is proportional to the number of genes. Major functional themes (immunity, development, transport, metabolism, signalling) are highlighted. Node color indicates transcriptional evidence from TCGA or METABRIC. **(G)** Selected immune signalling pathways with differentially expressed genes in IGX-positive breast cancers. Dot plots show interaction coefficients and significance values of pathway genes from IGX-positive samples in the two cohorts. **(H)** Association of IGX-positive luminal-A breast cancers with immunogenic characteristics in the TCGA cohort. Mann-Whitney U tests (FDR < 0.05).

To interpret this IGX functionally, we examined its genomic and immune features. For genomic features, 11q13.1 deletion associated with reduced expression of several genes at the deleted locus (FDR < 0.05; **Figure S4**). *MEN1*, the one established cancer gene in the region, was significantly downregulated in both TCGA and METABRIC (FDR < 10^-10^; **Figure 5C**). *MEN1* encodes menin, a tumor suppressor and transcriptional regulator involved in proliferation pathways [43]. *MEN1* is affected by somatic mutations or CNAs in multiple cancer types and its germline mutations predispose individuals to neuroendocrine cancers [44, 45]. For immune features, neutrophil levels in IGX-positive samples were consistently predicted as zero based in CIBERSORT, representing most luminal-A breast cancers overall (61% in TCGA and 71% in METABRIC; **Figure 5D**). Neutrophil levels alone were not associated with progression-free or relapse-free survival (**Figure 5A-B**). This may reflect complex anti-tumor and pro-tumor roles of neutrophils depending on their maturation and TME context [46]. A recent *in vivo* study linked *MEN1* activity in cancer to infiltration of neutrophils and CD8+ T cell [47]. These functions may interact synergistically with genomic losses of *MEN1* to drive tumor progression.

To explore the potential synergistic role of this IGX, we performed transcriptomic analyses to find differentially expressed genes in IGX-positive luminal-A breast cancers. In an integrative analysis of TCGA and METABRIC data, we found 227 differentially expressed genes with concordant differential expression in IGX-positive samples by controlling for the individual effects of 11q13.1 loss and reduced neutrophil levels (FDR < 0.1; **Figure 5E, Methods**). This included five known cancer genes (*ANK1*, *COL2A1*, *GPC3*, *KDM5A*, *TNFAIP3*). Integrative pathway enrichment analysis [48] highlighted biological processes consistently regulated in IGX-positive samples in the two cohorts, with major functional themes related to the immune system and inflammation, molecular transport, metabolism, signal transduction, and developmental processes (**Figure 5F**). Specifically, we observed pathway-level enrichments in interleukin signalling, STAT regulation, and neutrophil chemotaxis that were characterised by upregulation of several pathway genes (*IL6, IL6R, STAT3, TNF*) (**Figure 5G**). The IL6/JAK/STAT pathway is aberrantly upregulated in several cancer types including breast cancer, and known to drive tumor proliferation and invasiveness and suppress antitumor immunity [49], providing a potential functional interpretation of this IGX. Supporting this hypothesis, we observed that IGX-positive samples showed elevated levels of proliferation, HRD, aneuploidy, and increased expression of the ICI target *LAG3* (**Figure 5H**). Thus, the survival association of this IGX is confirmed in two independent cohorts of luminal-A breast cancer and the clinical phenotypes of these more aggressive cancers are explained by transcriptional deregulation of immune signalling pathways.

## DISCUSSION

Features of the tumor genome and its immune microenvironment jointly contribute to oncogenesis, disease progression and metastasis, and are therefore a major target of precision therapies and biomarkers. Using our new computational approach, we found 34 immunogenomic interactions in multiple cancer types in which the co-occurring genomic alterations and TME features may act synergistically, enhancing disease progression and leading to adverse outcomes. We also highlight the complexity of such genome-TME interactions by uncovering specific driver mutations and immune cell types that show either positive or negative associations with patient outcomes depending on the context of the genome and TME. Further associations with genome instability, cytolytic activity, and target expression of immune checkpoint inhibitor therapies provide complementary evidence and potentially actionable insights into IGXs.

We found that one IGX comprising the genetic deletion of the tumor suppressor *MEN1* combined with reduced neutrophils highlights a subset of luminal-A breast cancers with increased disease progression and relapse, as seen in two independent cohorts of breast cancers. Transcriptomic and pathway-level analyses suggest that IGX-positive luminal-A breast cancers show elevated levels of IL6/JAK/STAT signalling and immunogenic characteristics, providing potentially actionable insights into this subset of cancers and experimental hypotheses for follow-up work. Notably, a recent in-vivo study highlights novel oncogenic roles of *MEN1* in lung and pancreatic cancers via modulation of TME, indicating that *MEN1* knockout leads to reduced tumor growth and increased activity of CD8+ T cells and neutrophils [47]. Our observations from clinical cancer cohorts provide a complementary angle of tumor-immune interactions of *MEN1* and suggest that more work is needed to decipher this interaction in cancer models and clinically annotated datasets of cancer samples.

We developed the computational multi-omics analysis algorithm PACIFIC to systematically identify interactions between pairs of omics features that associate statistically with patient survival. We demonstrate this method by mapping IGXs comprising somatic alterations in cancer genomes and immune cell activities inferred from bulk cancer transcriptomes. However, the method is broadly applicable to other omics modalities, such as proteomics or epigenomics profiles. Besides patient survival information, other clinical or phenotypic information such as disease subtypes, grade or stage can be used to map informative multi-omics interactions. Overall, PACIFIC is a flexible computational method to integrate multi-omics profiles with clinical and phenotypic information to characterise the underlying biology and translational avenues.

Our study is subject to certain limitations. The genomic and immune features in the IGXs only represent a subset of potentially relevant elements of the tumor-immune interactome. We only considered a high-confidence set of frequently mutated cancer genes and recurrent CNAs, while other alterations such as structural variants or non-coding mutations were not considered. Future studies leveraging whole genome sequencing data will address this limitation. The signatures of major immune cell types derived from bulk transcriptome deconvolution do not capture the full complexity of TME and likely miss less-common and tissue-specific cell types. Recent advances in single-cell sequencing offer a vast increase in cellular resolution, however such datasets are currently limited in size and clinical breadth. IGXs highlight subsets of high-risk patients with certain genomic and TME characteristics, however these are not readily usable as biomarkers in clinical applications since IGXs need to be validated in additional cohorts and next, scalable low-throughput assays are required for translation. Lastly, our feature selection framework relies on generalised linear regression models that focus on linear relationships of multi-omics features. While these computational approaches are robust and easy to interpret, the assumption of linearity can limit their performance. Alternative models such as random forests and neural networks capture more complex associations, although these models often need more input data and provide only limited interpretability.

Our findings highlight an underexplored search space for mechanisms of disease progression, potential drug targets, and prognostic and predictive biomarkers. As single-cell RNA-seq approaches become applicable to larger clinical cohorts, we can create high-resolution maps of IGXs to deepen our understanding of the complex interactions of the TME of the tumor and its genomic alterations. Future work to develop advanced maps of IGXs along with experimental and clinical validations of these interactions will lead to mechanistic insights into cancer biology and enable advances in precision oncology.

## METHODS

### TCGA data collection

We retrieved data for 9201 primary, untreated cancer samples from 32 solid cancer types from the TCGA PanCanAtlas project [25] with matched genomic alterations, transcriptomic profiles, and clinical information. Each cancer type was analysed independently. We used either patient overall survival (OS) or progression-free survival (PFS) for cancer types based on prior recommendations [29]. Samples with missing survival information were excluded. Survival times were capped at 10 years with right censoring. We included patient age, patient sex, tumor stage, and tumor grade as baseline clinical variables. If stage and/or grade information was available for most samples then it was included in our analyses, and in those cases, samples with missing information were excluded. Genomic alterations included single nucleotide variants (SNVs) and insertion-deletions (indels) derived from whole-exome sequencing (WES), CNAs derived from Affymetrix SNP 6.0 (SNP6) arrays, and matching transcriptomics data from bulk RNA-seq experiments. We retained only cancer samples with three data categories available: (i) survival and baseline clinical variables, (ii) genomic alterations, (iii) gene expression data. For genomic alterations, we analysed small mutations (SNVs, indels) and CNAs separately as explained below. We split the breast cancer (BRCA) cohort into five subtypes based on PAM50 classification and split the lower-grade glioma (LGG) cohort into two subtypes based on mutational status of *IDH1* gene. We excluded cancer types with less than 100 samples and cancer types in which with survival events occurred in less than 5% of the cohort. Finally, our analysis included 8450 samples across 26 cancer types: *BLCA*, *BRCA-LumA*, *BRCA-LumB*, *BRCA-Basal*, *CESC*, *COAD*, *ESCA*, *GBM*, *HNSC*, *KIRC*, *KIRP*, *LGG-IDH^mut^*, *LIHC*, *LUAD*, *LUSC*, *OV*, *PAAD*, *PCPG*, *PRAD*, *READ*, *SARC*, *STAD*, *TGCT*, *THCA*, *THYM*, *UCEC* (**Table S1**).

### Driver mutations and recurrent CNAs

Two major types of genomic alterations were considered. First, we analysed 299 significantly mutated genes (SMGs) with somatic SNVs and indels derived previously from a TCGA PanCanAtlas publication [26]. Mutations with protein-coding impact were included and silent mutations were excluded. Second, we analysed recurrent CNAs comprising recurrently gained or lost genomic regions obtained from a previous TCGA PanCanAtlas publication [27] using the GISTIC2 method [50] (FDR < 0.25, confidence level 0.99). CNA data downloaded from the Firehose repository (https://gdac.broadinstitute.org; May 4th, 2020). CNA events for each cancer type were derived from the all_lesions output file of GISTIC2. The number of recurrent CNAs varied among cancer types, ranging from 18 CNAs in THYM to 99 CNAs in UCEC (**Table S1**). Recurrent CNAs and mutations in SMGs are referred to as genomic features and were defined as either present or absent per cancer sample.

### Immune cell levels

Each cancer sample was annotated based on the relative abundance of 22 major immune cell types predicted previously from bulk transcriptomes [18] using the CIBERSORT method [28]. CIBERSORT used the LM22 reference set of gene expression signatures for 22 types of immune cells. To provide robust and interpretable values of these predicted immune cell levels (ICLs), numeric values from CIBERSORT were grouped to ICL-high and ICL-low annotations using median dichotomisation separately for each cancer type and immune cell type. These annotations of cancer samples are referred to throughout the study as immune features. Median dichotomisation of immune features was performed separately in each iteration of feature selection procedure as described below.

### The PACIFIC method and finding immunogenomic interactions (IGX)

To discover survival-associated interactions of genomic and immune features, we adapted an elastic net framework from our previous studies [23, 24] to binary interactions of multiple omics datasets. Each cancer type was analysed separately. Recurrent CNAs and SMGs with small mutations were also analysed separately. Briefly, patient survival information was associated with IGXs by fitting regularised regression models on subsets of input data a series of 100,000 iterations. At each iteration, we selected a random 80% subset of patients to compile the iteration-specific input dataset (D) and used these to median-dichotomise ICLs into binary features over D. For a given genomic feature G and immune cell fraction ICL, each cancer sample was first assigned into one of four categories: [G-present, ICL-high], [G-present, ICL-low], [G-absent, ICL-high], [G-absent, ICL-low]. We applied three preprocessing steps. First, we removed sparse IGXs for which any of their four annotations spanned less than 5% of D. Second, we converted each IGX into a binary feature by selecting one of [G-present, ICL-high] or [G-present, ICL-low] as the main category, assuming that the presence of a genomic alteration rather than its absence is responsible for interactions with the immune system. The main feature (ICL-high or ICL-low) was selected based on a stronger survival association from univariate Cox proportional-hazards (CoxPH) models fitted to D (i.e., Wald test P-value). The remaining three categories were merged into a reference category, resulting in a binary annotation of each cancer sample [IGX-present *vs*. IGX-absent]. Third, we pre-filtered weak IGXs for which the survival associations were not significant (Wald P > 0.1). Next, to identify candidate prognostic IGXs at each iteration, we used CoxPH regression with elastic net regularisation [51]. A series of CoxPH models fitted on subsets of patient survival data and all potential IGXs were regularised. Model variables included preprocessed IGXs, common clinical variables (patient age, sex, tumor stage and/or grade; as available for cancer types), as well as the individual genomic and immune features contributing to the preprocessed IGXs. The elastic net models were applied using the R package *glmnet* (V4.1-1) [52] where the hyperparameter *λ*, which controls the strength of regularization, was tuned by five-fold cross-validation within each iteration, and the hyperparameter *α*, which controls the balance between L1 and L2 penalty, was fixed to 0.5. Regularised regression selected candidate IGXs from a larger space of potential IGXs. At each iteration, we recorded model variables with non-zero coefficients to reveal a subset of IGXs that were most associated with the survival data in that iteration. After completing all iterations, we identified candidate IGXs selected by regularisation in more than 50% of the total iterations. Next, candidate IGXs were filtered with four ANOVA tests. At this step and in all downstream analyses, the entire dataset rather than data subsets was used to median dichotomise ICLs to immune features. Each ANOVA test compared two nested CoxPH models: (i) null model including baseline variables and (ii) alternative model including the IGX and baseline variables. The four null models were: (i) CoxPH (Survival ∼ clinical variables), (ii) CoxPH (Survival ∼ clinical variables + G), (iii) CoxPH (Survival ∼ clinical variables + I), (iv) CoxPH (Survival ∼ clinical variables + G + I). These stringent comparisons were designed to select final IGXs based on two criteria: (a) providing complementary prognostic information to common clinical variables, and (b) providing complementary prognostic information to the genomic features (G) and immune features (I) alone. Collectively, IGXs passing these tests indicated interactions of genomic and immune features with respect to patient survival. P-values from the ANOVA tests were adjusted for multiple testing using the Benjamini-Hochberg false discovery rate (FDR) method [53]. Candidate IGXs that passed all four ANOVA tests (FDR < 0.05) were considered as the final prognostic IGXs. Univariate and multivariate CoxPH models were applied using the R package *survival* (V3.2-11).

### Interpreting IGXs with genomic features and immune features

To evaluate IGXs, we performed further survival analyses comparing each IGX with its components: the genomic features (*i.e.*, SMG mutations, CNAs), and the immune features. Survival analyses were performed separately for relevant cancer types. First, we compared survival rates by IGX features. Second, as controls, we compared survival rates by genomic features alone (G) and immune features alone (I). Multivariate CoxPH models (M1) with IGX, G, or I variables as well as clinical variables were fitted to the entire dataset of the corresponding cancer type. A null CoxPH model (M0) with only the clinical variables was also fitted as control. Hazard ratio (HR) values of features were obtained from M1, whereas P-values of features were calculated by ANOVA tests comparing M1 and M0. To further visualise IGXs, we ranked G and I features based on summed log-HR values across the IGXs, as well as the frequencies of G and I features across the IGXs. CNAs in the IGXs were manually annotated to identify the most likely known cancer gene as CNA targets, using the Cancer Gene Census (CGC) database [54]. First, we focused on the CGC genes overlapping with the cytobands corresponding to each CNA. Then we used non-parametric Mann-Whitney U tests to evaluate differential expression relative to the CNA events in corresponding cancer types. Significant genes (FDR < 0.05) were manually curated to select one representative gene for each CNA. Transcriptomics data was downloaded as TPM values from the Genomic Data Commons website (https://portal.gdc.cancer.gov; January 25th, 2023).

### Evaluating IGXs with measures of immunogenicity

We analysed IGXs using a comprehensive collection of genomic and immunological measures of cancers from a previous TCGA study [18]. The datasets were processed as follows. First, measures with missing values in more than 20% of all TCGA samples were removed. These included TIL regional fraction, BRC/TCR richness, BCR/TCR evenness, and BCR/TCR Shannon score. Second, non-silent and silent mutation rates were summed to derive a single measure of mutation rate per sample. Third, the genomic instability measures included the aneuploidy score while we excluded the number of genomic segments and fraction genome altered to reduce redundancy. Fourth, the indel neoantigen count was excluded as it was only available for a subset of TCGA samples. In addition to this collection, we also evaluated transcript abundance of four major immunotherapy targets (*PDCD1, CTLA4, CD274, LAG3*). We log-transformed the following measures to adapt these to linear regression analysis: transcript abundance (TPM values), cytolytic activity, somatic mutation rate, SNV neoantigen counts, homologous recombination defects, and aneuploidy score. We analysed these measures in the context of IGXs, using two related approaches. For each IGX, we first compared IGX-positive cancer samples with all other samples (i.e., IGX-negative samples) of the corresponding cancer type using non-parametric Mann-Whitney U tests and adjusted P-values for multiple testing using Benjamini-Hochberg correction at a significance level of 0.05. For a more stringent analysis, we selected the significant associations from the previous analysis and tested statistical interaction effects of IGX-positive samples relative to immune features alone (I) and genomic features alone (G). We used linear regression models with the immunogenomic features as response variables and IGX, as well as the respective G and I features as predictors. To select specific effects of IGXs in addition to its genomic and immune features alone, the significant interaction terms corresponding to IGXs were reported (P-value < 0.05; F-test) were reported.

### Validating the 11q13.1-neutrophil IGX in additional genomics cohort from METABRIC

One IGX comprising 11q13.1 losses combined with reduced neutrophil levels found in luminal-A breast cancer was validated in an independent cohort of primary luminal-A breast cancers from the METABRIC project [42]. The required datasets including CNAs, transcriptomics, and patient survival and clinical information were downloaded from cBioPortal (http://www.cbioportal.org; June 5th, 2024). We analysed the Discovery group of the METABRIC dataset. As CNA calls were available only for protein coding genes, we called 11q13.1 deletion events for samples in which more than 75% (48/64) of the genes located in 11q13.1 were deleted. The transcriptomics dataset included log2-transformed normalised intensities. We used the CIBERSORTx [55] web portal (https://cibersortx.stanford.edu) and with the LM22 gene signature matrix and default parameters to estimate the relative abundance of the 22 immune cell types. For survival analyses, we used relapse free survival and capped survival times at maximum of 10 years similar to our TCGA survival analyses. We fitted three multivariate CoxPH models to patient survival (M1) with one of IGX, G, or I as the predictor and the baseline clinical variables (patient age, tumor stage and grade) as covariates. We also fitted a control CoxPH model to patient survival (M0) with only baseline clinical variables included. P-values associated to each feature (IGX, G, or I) were derived ANOVA tests comparing models M1 and M0, whereas the log-HR value of each feature was derived from model M1. The IGX was validated in the Discovery subset of METABRIC while the Validation subset of cancer samples showed no significant associations.

### Transcriptomics analyses of 11q13.1-neutrophil IGX

RNA-seq data for the TCGA BRCA cohort was downloaded as raw counts and as TPM values from the Genomic Data Commons website (https://portal.gdc.cancer.gov, January 25th, 2023). Differential gene expression analysis was performed for TCGA BRCA-LumA and METABRIC-LumA cohorts using different approaches. TCGA RNA-seq data was processed with the DESeq2 method [56] with default parameters and raw counts as input. METABRIC microarray data was processed with the Limma method [57] with default parameters and log intensities as input. We used the design formula "expression ∼ IGX+G+I" with binary variables indicating the presence of the IGX feature and its genomic (G) and immune (I) features in each cancer sample. We extracted P-values and log2-transformed fold-change values of the IGX terms to capture the interaction between the genomic and immune features of the IGX. P-values were adjusted for multiple testing using the Benjamini-Hochberg method separately for each cohort. We also listed the genes overlapping with the genomic deletion at 11q13.1 and applied Mann-Whitney U tests to find differentially expressed genes responding to 11q13.1 deletion separately for TCGA and METABRIC datasets (FDR < 0.05).

### Pathway analyses of 11q13.1-neutrophil IGX

To functionally interpret the 11q13.1-neutrophil IGX in luminal-A breast cancer, we performed an integrative pathway enrichment analysis of its transcriptomic signatures from TCGA and METABRIC. We used the DPM method [48] to prioritise genes that were jointly either up-regulated or down-regulated in the two datasets based on the regression coefficients of the IGX term and corresponding P-values derived above. We defined the constraints vector in DPM as (1, 1). Molecular pathways from the Reactome database and biological processes from Gene Ontology were retrieved from the g:Profiler web server [58] (downloaded on June 7th, 2024) and gene sets with 25-250 genes were used in the analysis. To reduce potential biases, we used a custom gene background set that excluded the LM22 gene panel used in CIBERSORT as well as the genes not shared between the TCGA and METABRIC datasets. Significant pathways were selected (FDR < 0.01). A joint enrichment map visualisation showing the significantly enriched pathways from the two cohorts was created in Cytoscape using standard protocols [59]. Subnetworks were organised and annotated manually.

## Author Contributions

MBa: Conceptualization; Resources; Data curation; Software; Formal analysis; Validation; Investigation; Visualization; Methodology; Writing—original draft; Writing—review and editing. ZPK: Visualization; Methodology; Writing—review and editing. ATB: Visualization; Methodology; Writing—original draft; Writing—review and editing. MS: Validation; Visualization; Methodology. JT: Data curation; Methodology; Writing—original draft. KG: Methodology, Visualization; Writing—original draft. CM: Methodology; Writing—review. MB: Data curation; Writing—review. JR: Conceptualization; Data curation; Methodology; Resources; Supervision; Funding acquisition; Investigation; Writing—original draft; Project administration; Writing—review and editing.

## Acknowledgements

This work was supported by the New Investigator Award of the Terry Fox Research Institute (TFRI) to JR, Investigator Award to JR from the Ontario Institute for Cancer Research (OICR), the Discovery Grant of the Natural Sciences and Engineering Research Council (NSERC) (RGPIN-2023-04646), to J.R., and the Canadian Institutes of Health Research (CIHR) Project Grant (PJT-162410) to J.R. A.B. was supported by the Ontario Graduate Scholarship (OGS). M.Ba., K.C., and M.S. were partially supported by Medical Biophysics fellowships from University of Toronto. M.Ba. was also supported by a Caven Fellowship. D.P. was partially supported by an Ontario STAGE HostSeq fellowship. Funding to OICR is provided by the Government of Ontario. The results shown here are in whole or part based upon data generated by the TCGA Research Network: https://www.cancer.gov/tcga. This study makes use of data generated by the Molecular Taxonomy of Breast Cancer International Consortium.

