## Supplementary Figures for "Combinations of genomic alterations and immune microenvironmental features associate with patient survival in multiple cancer types"

Part (i)

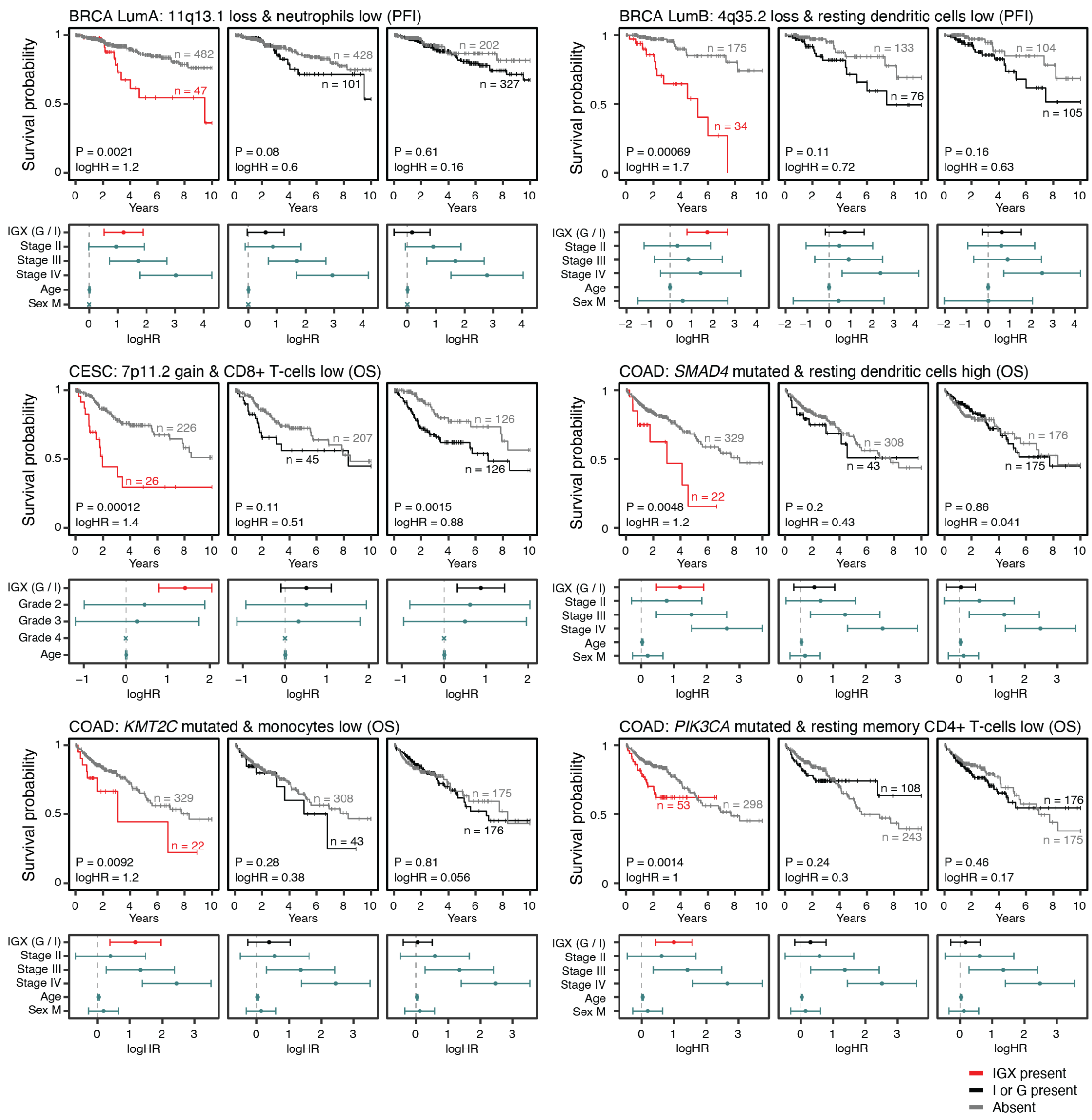


**Figure S1. Catalogue of IGXs with survival associations.** Kaplan-Meier (KM) plots (above) compare survival curves of IGX-positive and IGX-negative subsets of cancer patients. Overall survival (OS) or progression-free intervals (PFI) are shown for different cancer types. As controls, survival analyses of the corresponding genomic features (G) and immune features (I) are shown. Below the KM plots, HR values of variables included in the CoxPH models are shown with 95% confidence intervals. All P-values were derived from ANOVA tests of CoxPH models comparing survival associations of each feature (either IGX, G, or I) with the baseline clinical variables. All HR values were derived from CoxPH models with each feature (either IGX, G, or I) and the baseline clinical variables as covariates.

Part (ii)

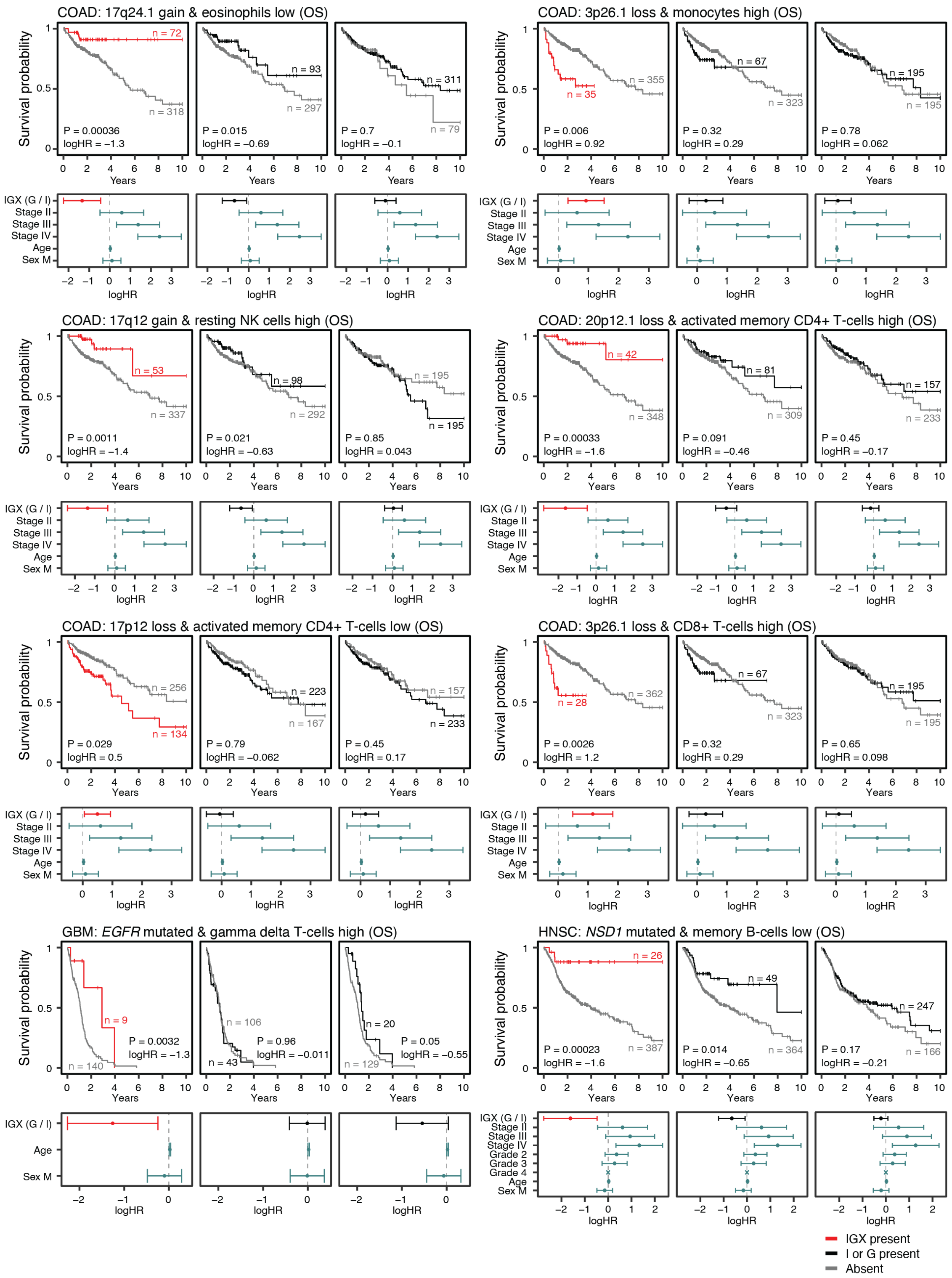


**Figure S1. Continued.**

Part (iii)

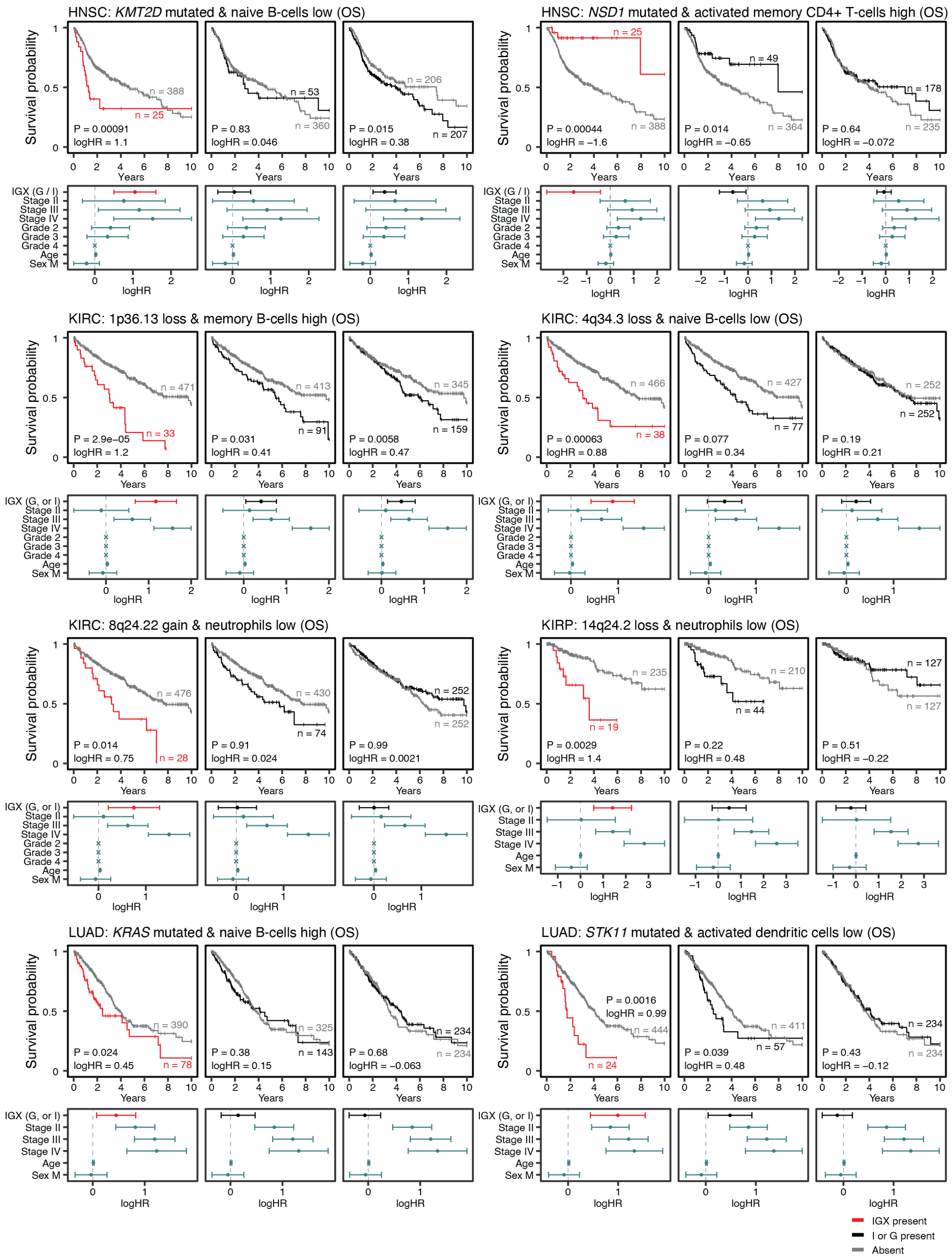


**Figure S1. Continued.**

Part (iv)

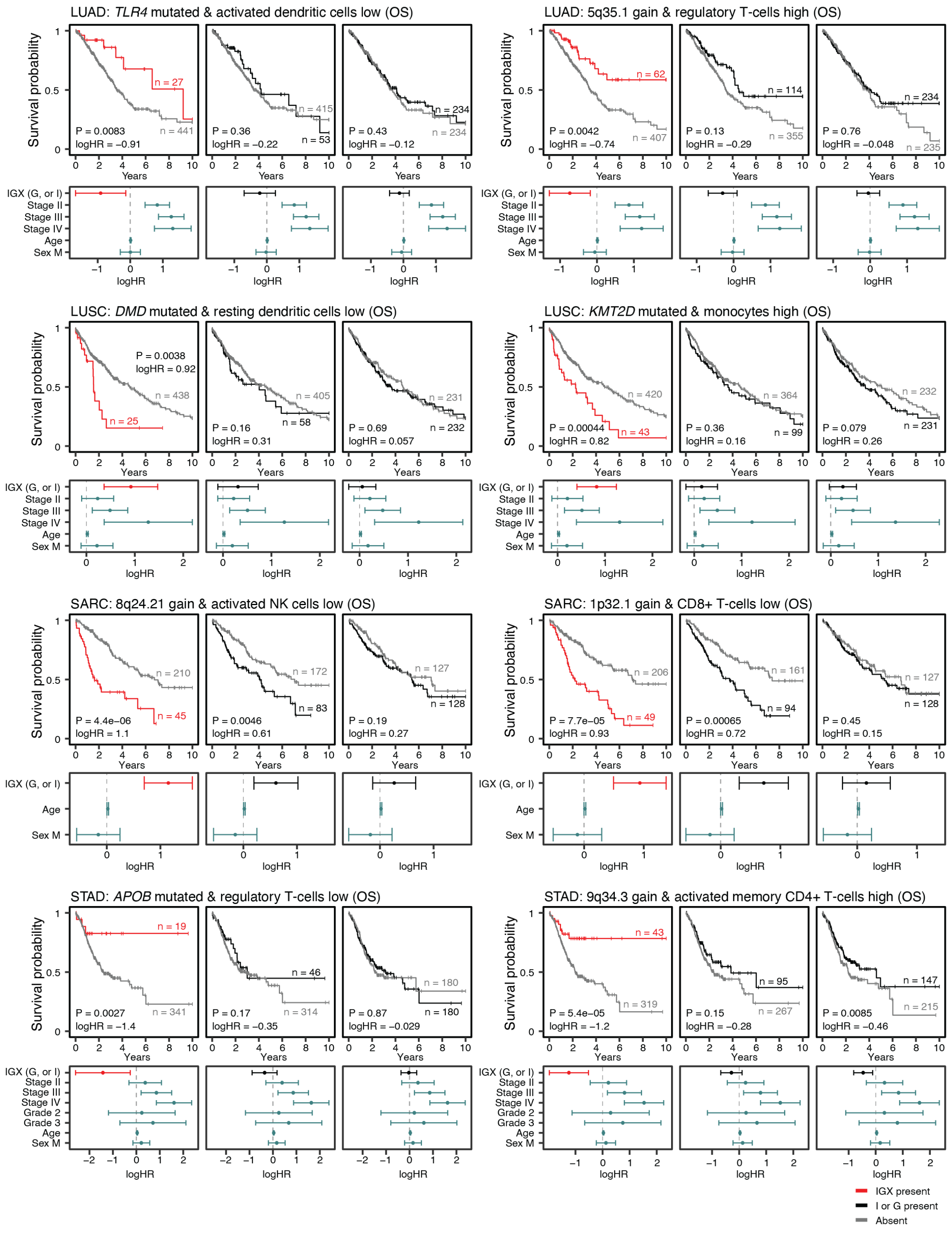


**Figure S1. Continued.**

Part (v)

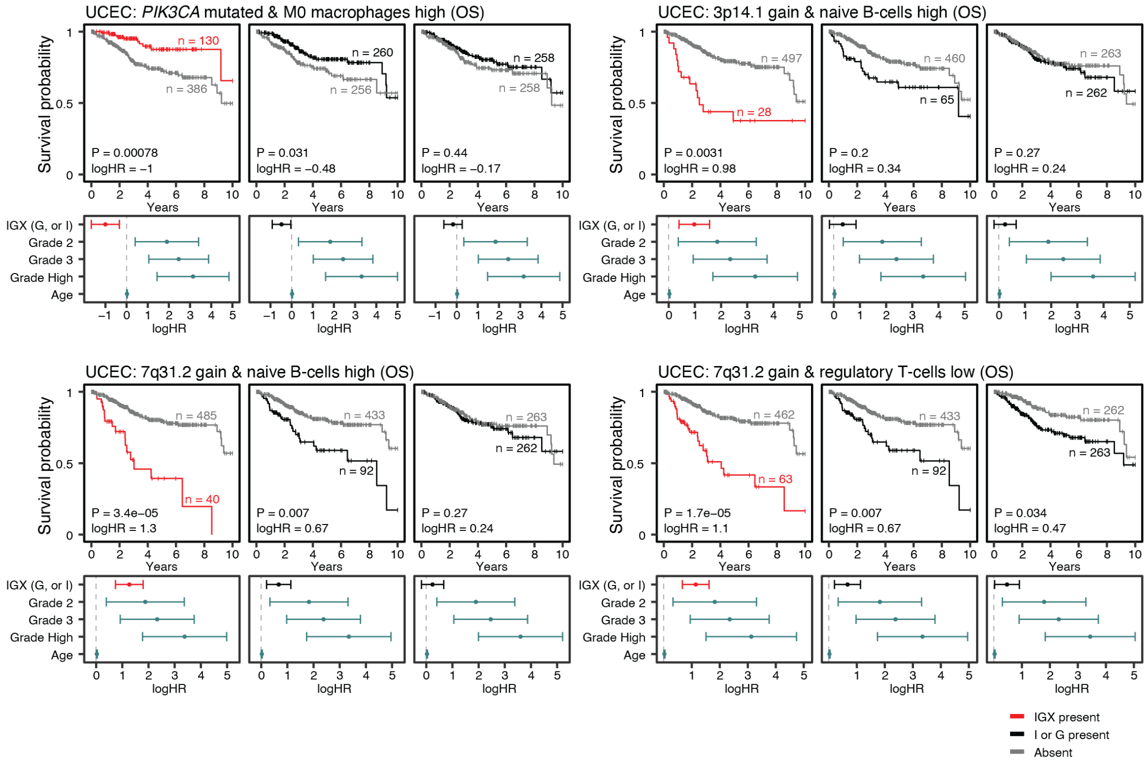


**Figure S1. Continued.**

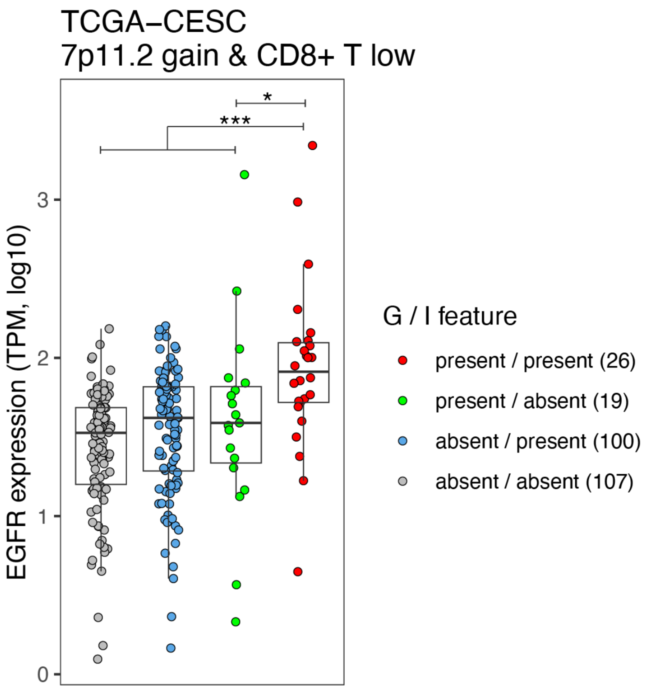


**Figure S2. *EGFR* expression in TCGA-CESC cohort.** Samples are grouped with respect to presence of 7p11.2 genomic gain and/or low CD8+ T status as defined in the corresponding IGX.

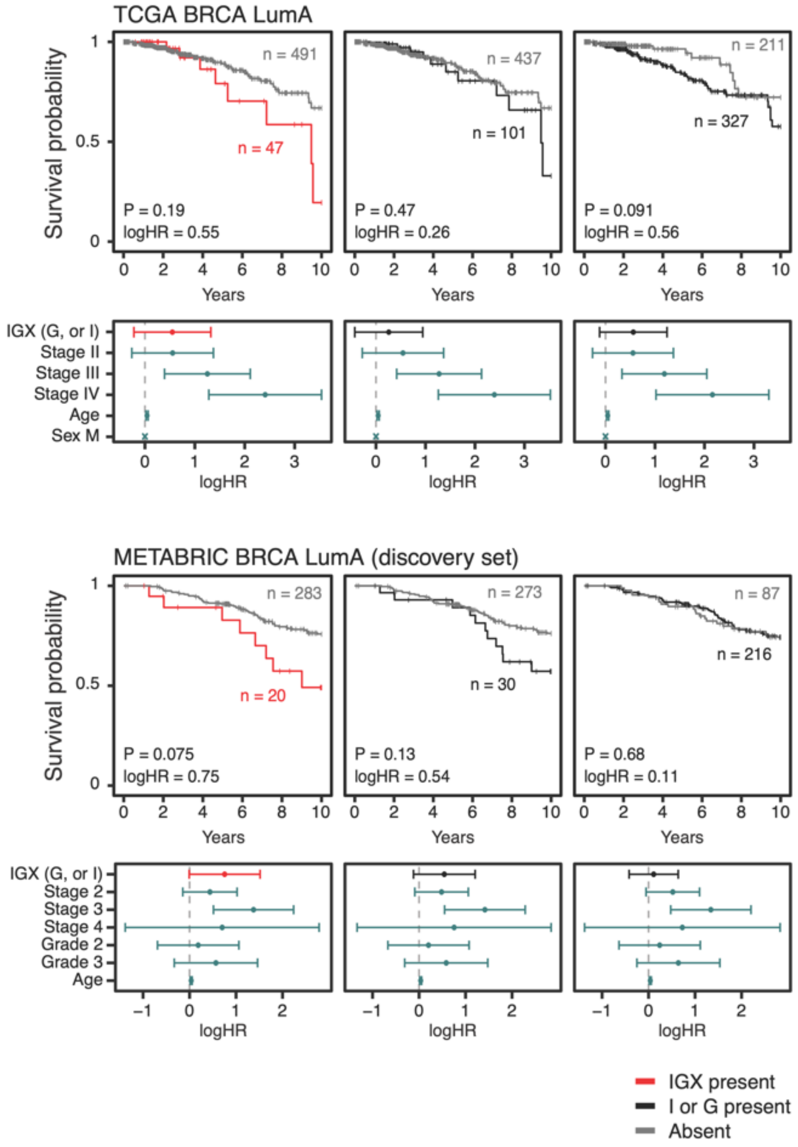


**Figure S3. Overall survival analysis of the IGX (11q13.1 loss, reduced neutrophils) found in luminal A breast cancer.** Top panel: TCGA cohort. Bottom panel: METABRIC (discovery set) cohort.

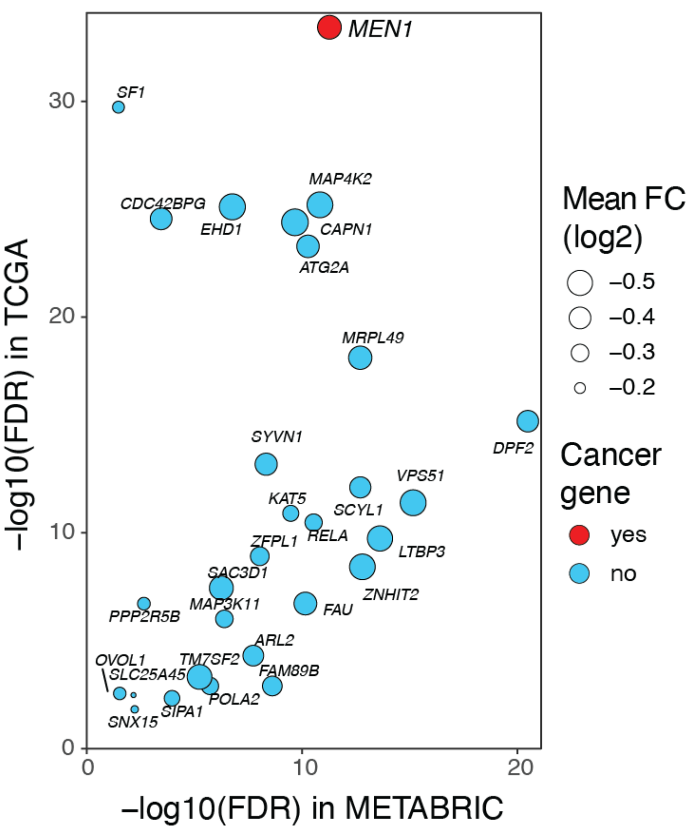


**Figure S4. Genes down-regulated in luminal-A breast cancers carrying the 11q13.1 deletion.** Genes in the 11q13.1 locus were separately analysed in TCGA and METABRIC cohorts by comparing their expression levels in 11q13.1-deleted and 11q13.1-balanced lumA-BRCA samples. The genes with consistent and significant signals in both cohorts are shown. The tumor suppressor *MEN1* is the only known cancer gene with highly significant down-regulation in both TCGA and METABRIC luminal-A samples according to the Cancer Gene Census database.
